# Food Supplementation Reduces Nematode Super-Shedding in a Wild Mammal

**DOI:** 10.64898/2026.01.16.699562

**Authors:** JSM Veitch, KE Wearing, J Mistrick, ME Craft, CE Cressler, RJ Hall, KM Forbes, SA Budischak

## Abstract

Anthropogenic changes to the environment, including altered food resource availability, influence host physiology, behaviour, and population dynamics, which can have strong downstream consequences on wildlife disease dynamics. Additionally, some individuals within a population contribute disproportionately to infection as ‘super-shedders’ of infection, but the extent to which food availability alters the likelihood of super-shedding and overall parasite infection patterns is poorly understood. We conducted a three-year field experiment in southern Finland to investigate how food supplementation and parasite removal affect nematode infection measures and the relationship between nematode infection and fitness of wild bank voles (*Clethrionomys glareolus*). Using a factorial design across 12 populations, we manipulated food availability and administered anthelmintic treatments to assess effects on nematode infection status, intensity, and two measures of super-shedding (abundance super-shedding, intensity super-shedding). We also examined parasite impacts on host fitness, including apparent survival probability and reproductive status. Food supplementation did not affect likelihood of infection, intensity or intensity super-shedding, but did reduce the likelihood of abundance super-shedding, suggesting an effect of food availability on infection heterogeneity. We also identified an interaction between nematode infection status and host age on fitness. Notably, infected younger individuals had reduced survival and reproduction, but infected older individuals had greater survival and reproduction compared to their uninfected counterparts. Our study provides novel empirical evidence on how anthropogenic changes in food availability can influence parasite transmission dynamics and the fitness consequences of these sub-lethal parasites in a wildlife system.

## Introduction

Anthropogenic environmental changes are increasingly linked to the emergence of infectious diseases (Gibb et al., 2024). Human modifications to food availability is one such feature that can have extensive impacts on wildlife (Birnie-Gauvin et al., 2017; Fehlmann et al., 2020). Changes in food availability and quality affect wildlife in terms of impacts on physiology (e.g. immunity, thermoregulation) (Fuller et al., 2021; Strandin et al., 2018), behaviour (Fuller et al., 2021; Newsome et al., 2015; Robb et al., 2008), population demographics (e.g. reproduction, survival, abundance) (Newsome et al., 2015; Robb et al., 2008), and ecology (e.g. distribution, disease dynamics) (Civitello et al., 2018; Robb et al., 2008). At the population level, greater food availability can increase host reproduction (Boutin, 1990; Nagy & Holmes, 2005) and survival rates (Briga et al., 2017; Martin, 1987), leading to higher density. Transmission of most pathogens increases with host density due to increased contact rates, so higher food availability may thus increase wildlife disease incidence (Becker et al., 2015).

At the individual level, increased food availability may instead reduce disease incidence (Becker & Hall, 2014). The association between resource availability and reduced infection risk has been extensively documented in laboratory experiments (e.g. Budischak et al., 2018; Clough et al., 2016; Coltherd et al., 2011; Israelson et al., 2024). Notably, changes in food availability can influence infection outcomes, including parasite fitness and harm of infection (Knutie et al., 2017) (Knutie et al., 2017). Knutie (2020) identified that food supplemented eastern bluebirds (*Sialia sialis*) had 75% less parasites than unsupplemented birds. However, increased food availability can elevate infection risk at the individual scale (Saino et al., 2000). Given the potentially contrasting effects of food availability between and within population and individual scales, the net effects of anthropogenic resource shifts on wildlife diseases are difficult to predict.

Field experiments that manipulate food sources can provide insight into the effects of food at both individual and population scales (Becker et al., 2015). A handful of food manipulation experiments in natural populations have found support that food can reduce parasite infections. For example, Agostini et al. (2017) found that increasing food availability in black capuchin monkeys (*Sapajus nigritus*) decreased helminth loads. Manipulating both food and parasite infection can also lead to further insights. Sweeny et al. (2021) found that experimental food supplementation reduced nematode infections, improved anthelmintic drug efficacy, and reduced likelihood of reinfection in wood mice (*Apodemus sylvaticus*). However, further research is needed to pinpoint the conditions where food supplementation can reduce or exacerbate infection risk, particularly those that manipulate both food availability and parasite infection.

In addition to reducing disease prevalence or intensity, alteration in food availability may affect how parasites are distributed among individuals within a population. Indeed, heterogeneity of parasite burdens is ubiquitous across many wildlife host-macroparasite systems (Shaw et al., 1998). Heterogeneity of infection can lead to a small fraction of hosts that disproportionately contribute to transmission due to high contact rates (i.e. super-spreaders) and/or infectiousness (i.e. super-shedders) (Chase-Topping et al., 2008; Kempf et al., 2022; VanderWaal & Ezenwa, 2016). Super-shedders may produce infectious parasites or pathogens at levels orders of magnitude greater than other infectious hosts and have consequently attracted considerable attention (Slater et al., 2016). However, the effect of food resources on infectious heterogeneity has not been widely studied. Notably, food quality has been shown to increase uneven distributions for *Daphnia* parasites (Vale et al., 2013) and some lab mouse parasites, but not others (Budischak et al., 2015). Alternatively, food supplementation may allow all individuals better access to adequate food resources for parasite defence (Sweeny et al., 2021), and consequently reduce super-shedders. Despite the potential for food to greatly alter parasite infection heterogeneity, these patterns remain unexplored in wild populations.

In recent decades, there has been growing recognition of the importance of studying sub-lethal parasites (Koltz et al., 2022; Shanebeck et al., 2022). The ecological consequences of sub-lethal parasites in natural systems are likely widespread but overlooked (Koltz et al., 2022). While sub-lethal parasite infections may not be consistently associated with pathological signs or mortality, these parasites can significantly influence host energetics and consequently host fitness (Shanebeck et al., 2022). Notably, sub-lethal parasites can exert negative effects on host reproduction and survival, but also host feeding rates, body mass, and body condition (Koltz et al., 2022). Given that these fitness-related traits are closely tied to food availability, experimental approaches can be a powerful tool to examine effects of sub-lethal parasites on hosts under different nutritional contexts.

We conducted a population-scale food and parasite manipulation experiment to investigate the interaction between food and helminth infection in wild bank voles (*Clethrionomys glareolus*). Examining sub-lethal helminth infections, and heterogeneity in infection, in bank voles is crucial, as infection burden could predict coinfection with zoonotic pathogens such as Puumala hantavirus (Guivier et al., 2014; Salvador et al., 2011). We investigated three key questions: (1) What is the effect of food availability on parasite infection status and intensity? (2) Does food availability affect the heterogeneity of parasite infections, specifically the likelihood of super-shedders? (3) How does food availability affect the fitness costs of sub-lethal parasite infections? We predict that food supplementation will lead to decreases in measures of parasite infection, as seen in other wildlife systems (Agostini et al., 2017; Knutie, 2020; Sweeny et al., 2021). We also predict that food supplementation will increase the likelihood of an individual becoming a super-shedder (Budischak et al., 2015). Lastly, food supplementation may reduce fitness costs associated with sub-lethal parasite infections.

## Materials and Methods

### Field Methods

We conducted a 3-year field experiment in southern Finland to test the individual and interactive effects of food availability and nematode infection on nematode infection dynamics, associated fitness costs, and super-shedder status in wild bank voles (*Clethrionomys glareolus*). The study was conducted across twelve populations that were randomized across four factorial treatment combinations of food supplementation and anthelmintic treatment: 1) fed-deworm, 2) unfed (i.e. not supplemented)-deworm, 3) fed-control (i.e. no anthelmintic treatment), and 4) unfed-control (i.e. no manipulations). Each treatment combination was replicated at three forest population sites that were at least 2 km apart to prevent movement of individuals between populations. Each site consisted of a 100m by 100m (1 ha) trapping grid with 61 evenly spaced Ugglan multi-capture live traps (Grahnab, Sweden) 10m apart. Two sites were replaced (using the same treatment combinations) over the winter periods between trapping years, either due to logging or consistently low vole numbers. This resulted in a total of 14 sites.

The food supplementation was provided from May through mid-November each year, where fed sites received food supplementation every two weeks. Following a similar protocol to Sweeny et al. (2021), the food mix for each site consisted of 7.875 kg of sunflower seeds (6350 kcal/kg) and 7.875 kg of mouse chow pellets (Altromin 1324 [3227 kcal/kg]; Altromin Spezialfutter GmbH & Co. KG, Lage, Germany). Unfed sites received no food supplementation. Under the anthelmintic treatment, captured voles in the deworm sites were treated once per month with an oral dose of a combined anthelmintic: ivermectin (10mg/kg; Knowles et al., 2013) and pyrantel (100mg/kg; Ferrari et al., 2009). This dosage is known to be effective at reducing adult and larval nematode loads (Sweeny et al., 2021). Nematode species present in these populations include *Heligmosomoides glareoli, Heligmosomum mixtum* and, less commonly, a Capillarid species (Chan & Budischak, unpublished data; prevalence = 0.011). Voles in the control groups received an equivalent weight-based volume of ~10% sugar water as a placebo.

Voles were longitudinally monitored from May through October each year when there is no snow layer to prevent trapping, following methods detailed in Mistrick et al. (2024). Trapping occasions occurred monthly, each with four checks over two consecutive days. At a vole’s initial capture, we fitted a Passive Integrated Transponder (PIT) tag (‘Skinny’ PIT tag, Oregon RFID, USA) for unique identification. Treatment (anthelmintic or sugar water) was administered only on the first capture of an individual within each monthly trapping occasion. A small proportion of non-reproductive voles (mean 2.4% ± 0.49% of the population) were culled at each site for immunological assays in August-September 2021, July-August 2022, and July 2023 (data not reported here) to minimize impacts on vole population dynamics. We calculated the number of voles captured per site and occasion divided by the site area (1 ha) as a measure of population density. Only the first capture per individual was included in the dataset for each month, ensuring that repeated measures reflect monthly intervals rather than multiple captures within the same month.

For the first capture of an individual each month, we recorded an individual’s sex, reproductive status, and head width (see Mistrick et al. (2024) for details). Head width, measured with calipers, served as a proxy for age (Kallio et al., 2014). Body condition was calculated as the residuals from a linear mixed-effects model (‘lme4’ package version 1.1-37; Bates et al., 2025). with log-transformed body mass as the response variable, log-transformed head width as a fixed effect, and handler as a random effect. This approach provides a body condition index by correcting for structural body size from body mass, following recommendations for using mass– size residuals as a condition metric (Schulte-Hostedde et al., 2005). We excluded pregnant females from this calculation.

Fecal samples were collected at each trapping occasion and traps were cleaned and replaced after each capture to prevent contamination. Fecal samples were subsequently processed using a salt flotation method (Pritchard & Kruse, 1982) to quantify nematode infection status (binary variable; 0 = uninfected, 1 = infected) and intensity, measured as eggs per gram of feces (EPG) among infected individuals (Margolis et al., 1982), which is correlated with adult worm count in these populations (Budischak and Veitch, unpublished data). We excluded fecal samples weighing less than 0.01g since they are less reliable measures of infection status and intensity (Budischak and Veitch, unpublished data). Three nematode species are present in these populations: *Heligmosomoides glareoli, Heligmosomum mixtum*, and less commonly a Capillarid species (0.011 prevalence; Chan & Budischak, unpublished data). Since the deworming medication is expected to reduce all three nematode species, and eggs from *H. glareoli* and *H. mixum* cannot be distinguished morphologically, infection status and intensity measures represent combined infections across all three nematode species.

### Statistical Analysis

To examine patterns in nematode infection status and intensity, we used generalized linear mixed-effects models (‘glmmTMB’ package version 1.1.11; Brooks et al., 2017). We used the simulateResiduals() function from the ‘DHARMa’ package (version 0.4.7; Hartig, 2020), which simulates residuals from fitted models, to evaluate goodness-of-fit and detect potential deviations from model assumptions. We first investigated nematode infection at the population scale by modelling population-level nematode prevalence as a function of our experimental treatments (food supplementation, anthelmintic treatment) and year, using a binomial distribution and including site as the only random effect. Prevalence was calculated as the proportion of infected individuals of all captured individuals at each site per month.

At the individual scale, we modelled nematode infection status as a binary variable (0 = uninfected, 1 = infected) using a binomial distribution. We modelled nematode intensity using a Gamma distribution with a log link. Since feces were deposited in traps prior to handling and anthelmintic treatment administration, all fecal samples at first capture represent pre-treatment infection levels. Thus, the pre/post-treatment variable distinguishes between an individual’s first capture (prior to any anthelmintic treatment effects) and subsequent monthly captures, after the anthelmintic (or sugar water) had the potential to affect nematode infection. Trapping occasion (measure of month of capture) was included as a second-order polynomial to capture non-linear seasonal trends in our response variables. The only exception to including trapping occasion as a second-order polynomial was in models limited to the first two months of data, where linear occasion was used instead. Predictors for both the infection status and intensity models included the main and interactive effects of food supplementation treatment (fed/unfed) and anthelmintic treatment (deworm/control), the main and interactive effects of anthelmintic treatment and pre/post-treatment, and occasion, year, vole sex, age, body condition, and population density. Models with recaptures (i.e. models with full dataset that were not subsetted to the first two months) included tag (vole individual ID) as a random effect to account for repeated measures. In addition to running these analyses on the full dataset, we ran these analyses on a subset of only captures in the first two months of the study to assess baseline infection levels before treatment. The first two months were selected because there were insufficient vole numbers in May 2021 alone. For any post-hoc comparisons of interactions, we used estimated marginal means (EMMs) with the ‘emmeans’ package (version 1.11.1; Lenth, 2023) to assess pairwise differences. Specifically, we applied the emmeans() function to extract marginal means from the fitted model, followed by Tukey-adjusted pairwise comparisons using the pairs() function.

We tested if food and/or anthelmintic treatment affected the distribution of infections among individuals, and highly infectious individuals (“super-shedders”) in particular. Defining supershedding is challenging and various metrics have been used depending on the host pathogen system (Kempf et al., 2022). Following the “80-20 rule”, captures with a nematode EPG in the top 20% of all infected samples (≥ 181.5 EPG) were classified as intensity super-shedders and captures with a nematode EPG in the top 20% of all samples (≥ 35.5 EPG) were classified as abundance super-shedders. Intensity super-shedders measure heterogeneity in parasite burden or infectiousness. In contrast, abundance super-shedders combine both likelihood of being infected and parasite burden, which we do not capture in any of our other infection metrics. Intensity super-shedders are primarily examined in the context of mathematical models (as a component of transmission (Lloyd-Smith et al., 2005) and in lab experiments for comparisons among infected individuals after a controlled exposure (Budischak et al., 2015; Dolinski et al., 2022; Gopinath et al., 2013), but have also been applied to assess supershedding in livestock (Courcoul et al., 2011; Matthews et al., 2006; Slater et al., 2016) and in wildlife (Santos et al., 2015). Measuring abundance super-shedders is of additional importance, as it captures the variation in transmission potential across the entire population, not just among infected individuals (e.g., Cooper et al., 2019). This created two binary variables indicating whether a vole was a (intensity, abundance) super-shedder for each capture. To assess the effect of food and anthelmintic treatments, and their interaction, on nematode intensity and abundance super-shedding, we used generalized linear mixed-effects models with binomial distributions. Vole individual ID was included as a random effect.

Lastly, we constructed individual-scale models to investigate the factors affecting two fitness measures - apparent survival and reproductive status. Apparent survival was modelled with a binomial distribution (0 = final capture, 1= subsequently captured). Predictors included the main and interactive effects of nematode infection status and vole age, as well as anthelmintic treatment, vole sex, reproductive status, body condition, and population density. This model excluded voles that tested positive for Puumala hantavirus to avoid any potential virus-induced mortality. For reproductive status (binomial distribution), predictors included the main and interactive effects of anthelmintic treatment and pre/post-treatment, and the main and interactive effects of nematode infection status and vole age. The data for this model was subsetted to voles ≥11.3 mm head width to include only voles within the age range where at least one individual was identified as reproductively active.

Plots were constructed using ‘ggplot2’ package (version 4.0.0; Wickham, 2011). For the interaction plots, partial residuals were extracted using ‘visreg’ package (version 2.7.0; Breheny & Burchett, 2017) and then also constructed using ggplot2.

## Results

After we excluded captures without fecal egg count data (e.g. no feces collected, fecal sample < 0.01g), we had a final dataset of 2,524 captures of 1,967 individuals, ranging from 1 to 9 captures per individual across different monthly trapping occasions (mean = 1.265 captures per individual). The overall prevalence of nematode infection was 38.5%, and the average nematode abundance was 54.01 ± 8.844 eggs per gram (EPG).

Anthelmintic treatment significantly reduced nematode prevalence across the populations (p = 0.003; Table S1, Fig S1), confirming its efficacy at the population scale. To further validate the efficacy of the anthelmintic treatment, we compared infection levels in the deworm and control sites in the first two months of the study to the overall study period (over 3 years). As expected, we observed no initial differences in nematode infection status and intensity across any of the treatment groups in the first two months of the experiment (infection status: food supplementation treatment (p = 0.333), anthelmintic treatment (p = 0.641), interaction (p = 0.678), intensity: food supplementation treatment (p = 0.795), anthelmintic treatment (p = 0.800), interaction (p = 0.674); Table S5). Importantly, when examining the effect of parasite removal at the individual scale, we found that the anthelmintic treatment interacted with time (pre/post-treatment) to significantly reduce the likelihood of nematode infection (infection status: p = 0.008) and a marginally significant reduction in nematode intensity (p = 0.063) (Table S2, Fig S4). Post-hoc comparisons revealed that individuals in the dewormed sites were significantly less likely to be infected post-treatment than control individuals (p < 0.001). Anthelmintic treatment also reduced the likelihood of an individual becoming a nematode egg abundance super-shedder (p < 0.001; Table S3, Fig 1), but not intensity super-shedder (p = 0.648; Table S3, Fig S3).

**Figure 1.**
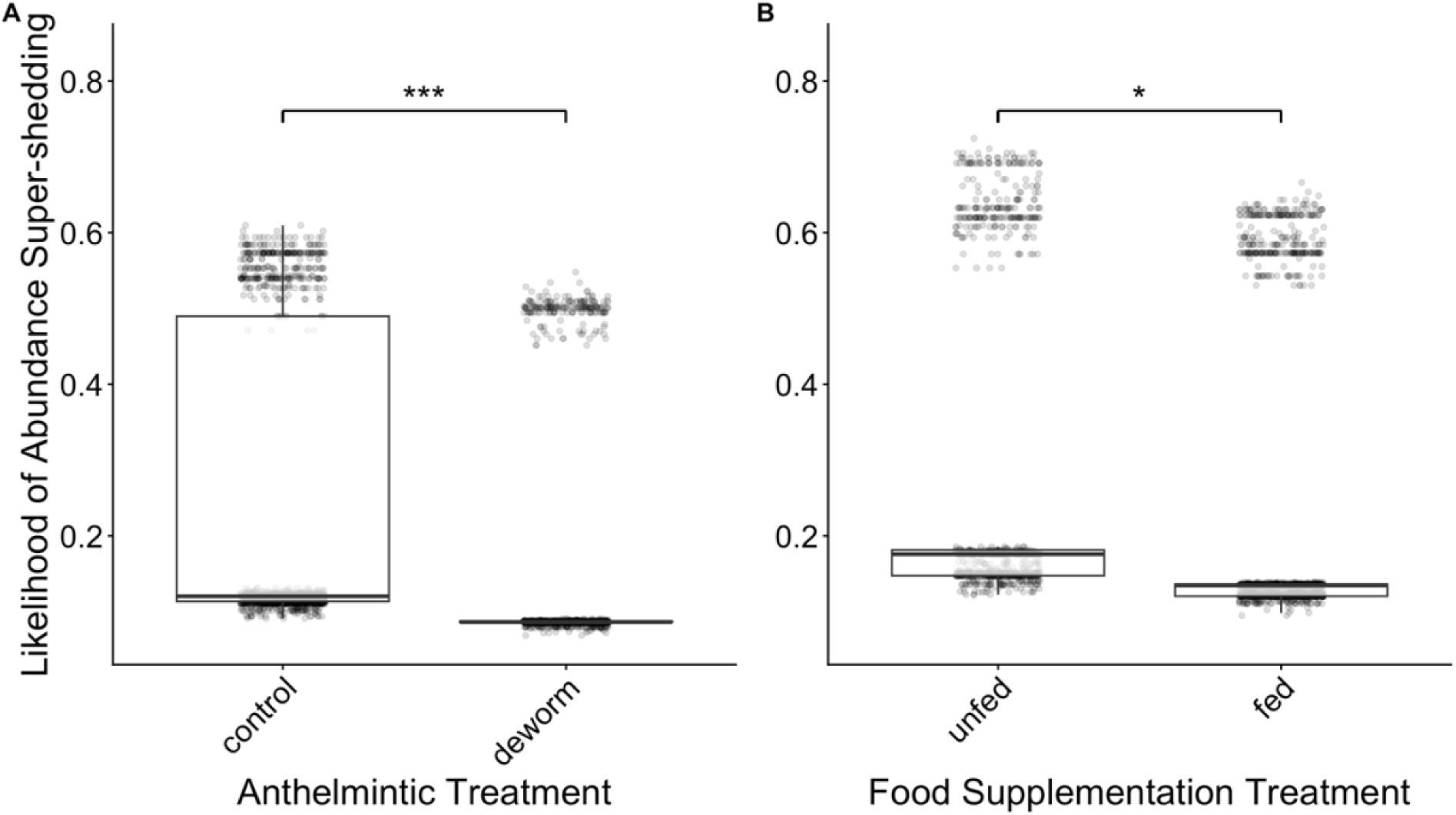
Bar plots displaying the effect of (A) treated anthelmintic and (B) food supplementation treatment on the proportion of nematode egg abundance super-shedder captures (top 20% of nematode EPG values from all captures; n = 2524 captures, 1707 individuals). Points in both panels represent partial residuals. Asterisks indicate significant differences between groups. treated anthelmintic and food supplementation both reduced the likelihood of an individual becoming a nematode egg super-shedder.

We found that food supplementation did not significantly affect population-level nematode prevalence (p = 0.091; Table S1, Fig S1). At the individual level, food supplementation had no significant effect on the likelihood of infection (p = 0.360), infection intensity (p = 0.373; Table S2, Fig S2), or likelihood of intensity super-shedding (p = 0.128; Table S3, Fig S3). However, we identified that food supplementation did reduce the likelihood of abundance super-shedding (p = 0.021; Table S3, Fig 1). This effect remained consistent regardless of anthelmintic treatment, as we found no significant interaction between food supplementation and anthelmintic treatment on super-shedding (p = 0.228; Table S3).

We identified several additional factors that shaped infection patterns at the individual level (Table S2, Fig S5). Infection status followed a shallow U-shaped seasonal pattern, with likelihood of infection decreasing from spring to autumn (p < 0.001) and infection intensity peaking in October (p = 0.016). Infection status varied across years (p < 0.001) and increased with age (p < 0.001); however, infection intensity decreased with age (p = 0.011). Thus, older individuals were more likely to be infected but carried lower nematode burdens. We also found that individuals with higher body condition were more likely to be infected (p = 0.022). Additionally, population density was negatively correlated with likelihood of infection and infection intensity, where individuals in higher-density populations were less likely to be infected (p = 0.017) and infected individuals harboured lower nematode burdens (p = 0.011). From these findings, seasonal, demographic, and ecological factors may have a more important relationship with infection patterns compared to food supplementation.

Although anthelmintic treatment did not affect survival or reproduction, infection status was correlated with both, but in an age-dependent manner (Table S4). Specifically, infection intensified age-related effects, such that young, infected voles had even lower apparent survival (p = 0.007; Table S4, Fig 2) and were less likely to be reproductively active (p = 0.007; Table S4, Fig 3) compared to uninfected young voles. Alternatively, older infected voles had greater apparent survival and likelihood of being reproductively active than their uninfected counterparts. While Anthelmintic treatment did not directly influence vole fitness, nematode infection itself may still play a significant, age-dependent role.

**Figure 2.**
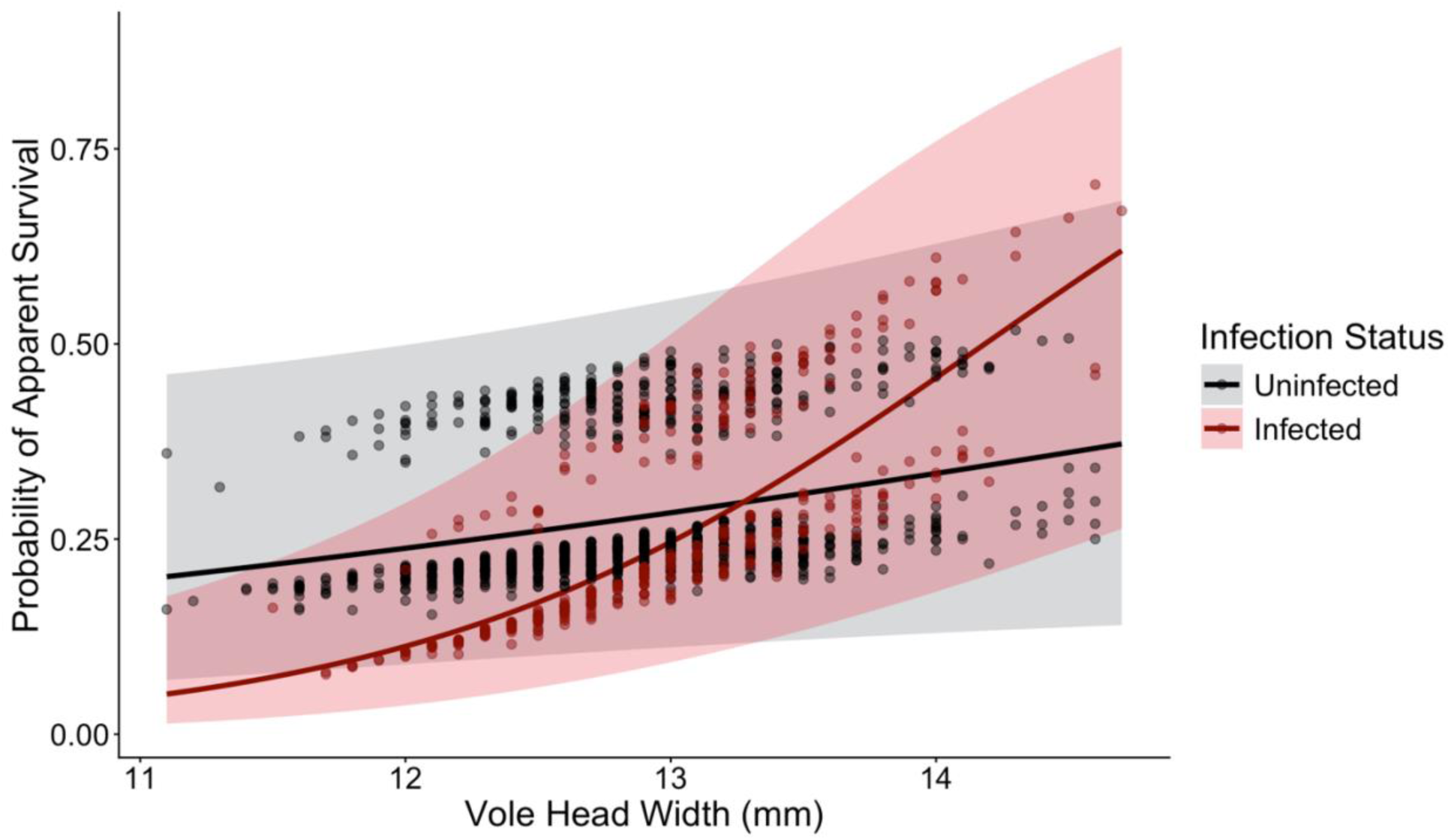
Fitted line plot of apparent survival probability across age (measured as head width in mm) for individuals infected with nematodes and uninfected. Lines show fitted values from model in Table 4 with shaded 95% confidence intervals. Points represent partial residuals. Individuals that tested positive for Puumala hantavirus were excluded from this dataset. Infected young voles had lower apparent survival and infected old voles had greater apparent survival compared to their uninfected counterparts (n = 1244 captures, 947 individuals).

**Figure 3.**
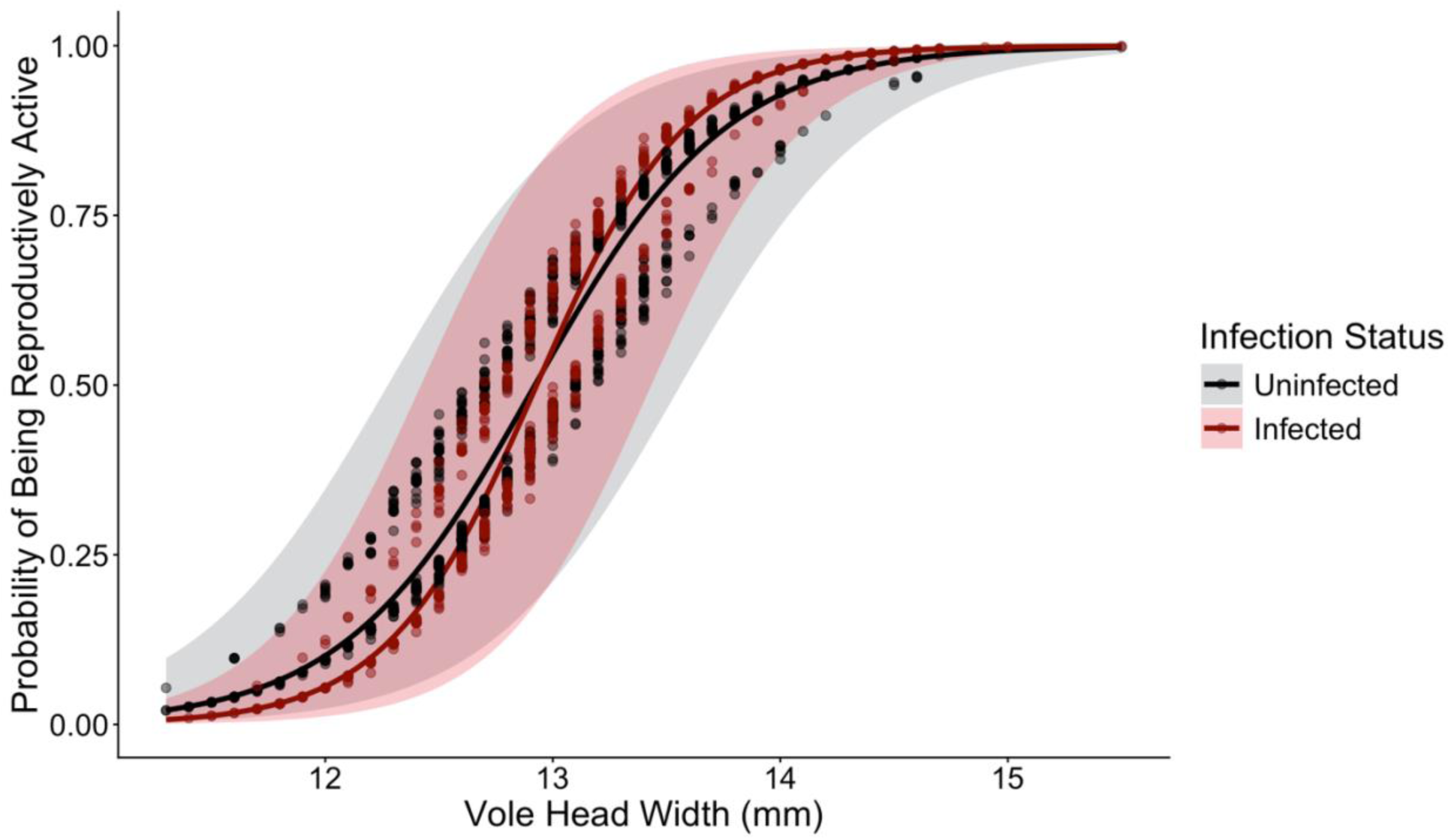
Fitted line plot of probability of being reproductively active across age (measured as head width in mm) for individuals infected with nematodes and uninfected. Lines show fitted values from model in Table 4 with shaded 95% confidence intervals. Points represent partial residuals. Analysis restricted to voles ≥11.3 mm head width to include only voles within the age range where at least one individual was identified as reproductively active. Infected young voles had slightly lower probability of being reproductively active and infected old voles had slightly higher probability of being reproductively active compared to their uninfected counterparts (n = 2446 captures, 1676 individuals).

## Discussion

Anthropogenic food provisioning is known to alter parasite infection outcomes and transmission in wildlife systems (Becker et al., 2015). However, the effect of varying food availability on parasite infection patterns at population and individual scales, including heterogeneity of infection, is poorly understood. We tested how food supplementation and anthelmintic treatment may interact to affect nematode infection patterns and the subsequent consequences of these infections on host fitness outcomes. Surprisingly, food supplementation did not alter nematode prevalence, likelihood of infection, intensity, or likelihood of intensity super-shedding. In contrast to these results, food supplementation did significantly reduce the likelihood of abundance super-shedding. Therefore, food availability may alter parasite transmission potential, even when the overall infection presence and burden exhibit no change, which holds implications for both parasite and wildlife health management programs.

The effects of food supplementation on super-shedders are particularly important, as they disproportionately drive transmission and hold management implications (Kempf et al., 2022). In our study, food supplementation reduced the likelihood of an individual to be an abundance super-shedder, but not nematode prevalence, likelihood of infection, intensity, or intensity super-shedder status. The effect of food supplementation on abundance, but not intensity, super-shedder status suggests that considering both the probability of being infected and parasite burden may be important in how food influences infection heterogeneity. Abundance super-shedding status is the only measure to consider the measure of nematode infection in the full host distribution (Rozsa et al., 2000), and therefore the only measure to respond to changes in either process simultaneously. It may be that while food supplementation does not influence solely nematode infection status or intensity, it does alter an integrated metric that considers both processes. The lack of a food effect on these other infection measures may also be because the cost of fully eliminating parasites may be greater than living with the infection (Viney et al., 2005) or that the continual presence of infective parasite life stages in the environment may lead to repeated exposure, as seen in other systems (Carlsson et al., 2012; Craig et al., 2009; Knowles et al., 2013; Müller-Klein et al., 2019). While changes in overall infection patterns may be limited, this does not preclude effects on extreme phenotypes across a full host population, such as abundance super-shedders.

Given that food acquisition can vary between individuals, especially in food hoarding species (Bohn, 2023), food supplementation may dampen differences in nutritional intake across a host population. Modelling work suggests food supplementation can reduce the proportion of super-shedders in a population by homogenizing resource allocation to immune defence (Hall, 2019). Additionally, field voles (*Microtus agrestis*) with high food availability had stronger immune responses against nematode infections (Forbes et al., 2016). Thus, food supplementation may disproportionately benefit individuals under the highest nutritional stress, improving their immune responses and reducing the risk of super-shedders emerging. Conversely, high protein diets increased likelihood of super-shedding of one helminth species, but led to a negligible decrease in another species in lab mice (Budischak et al., 2015). Food supplementation may not always reduce super-shedding, highlighting the need to study nutritional effects across diverse host-parasite systems, especially in natural settings where greater environmental and individual variation may alter infection heterogeneity. Our result suggests nutritional interventions could mitigate transmission risk in wildlife, especially when targeted towards food-limited hosts.

Another important finding from our study is that nematode infection altered fitness outcomes depending on host age. While apparent survival probability generally increased with age, we identified nuanced differences depending on whether individuals were infected. Notably, infected younger voles had lower apparent survival probability than uninfected younger voles. In contrast, infected older voles had greater survival than their uninfected counterparts. These contrasting results may be due to age-related differences in infection responses. Juveniles are often immunologically naive and may also experience energetic constraints from trade-offs between parasite defence and growth or future reproduction (Ashby & Bruns, 2018; Garrido et al., 2016). These limitations may leave younger individuals more vulnerable to parasite-related mortality. In contrast, older individuals may have developed tolerance to infection (Jackson et al., 2014). Older voles may be those that can persist with nematode infections and are more likely to reach adulthood. This aligns with another finding from our study that there was a positive correlation between body condition and likelihood of infection. Older individuals may also benefit from prior nematode exposure that could strengthen their immune response against bacterial or viral pathogens (Chen et al., 2023). Any or a combination of these mechanisms could potentially enhance survival of infected individuals later in life by enhancing their ability to cope with or fight off future challenges.

We found a similar result in how nematode infection interacts with age to alter the likelihood of a host to be reproductively active. Younger infected voles were less likely to be reproductively active than their uninfected counterparts whereas older infected voles were more likely to be reproductively active. Analogous to our previous explanation of juvenile vulnerability to parasitism in terms of survival, immunological naivete and energetic constraints could lead to reduced reproduction in response to nematode infection for younger voles (Ashby & Bruns, 2018; Garrido et al., 2016). Similarly, older individuals may have developed sufficient immune responses to tolerate infection, allowing them to survive and reproduce despite being infected (Jackson et al., 2014). An additional explanation for why older infected individuals may have a greater likelihood of being reproductively active is that reproductive activity itself may increase parasite susceptibility, either due to changes in energy allocation or immunomodulation by reproductive hormones. Energetic trade-offs between reproduction and immune defence could leave reproductively active individuals more vulnerable to infection. For instance, the cost of lactation in red deer (*Cervus elaphus*) (Albery et al., 2021) and number of offspring to care for in female collared flycatchers (*Ficedula albicollis*) (Nordling et al., 1998) has been linked to greater parasite burdens. Additionally, increases in testosterone in males are closely tied to reproduction and can suppress the immune response and alter infection (Folstad & Karter, 1992; Husak et al., 2021). Therefore, differences in reproductive activity between infected and uninfected voles may be due to infection reducing reproduction in younger voles but also increased tolerance or susceptibility in older voles.

This experimental field study demonstrated the effect of food supplementation on heterogeneity of parasite transmission in a wildlife system. Surprisingly, in contrast to multiple theoretical studies (Becker & Hall, 2014; Erazo et al., 2022), food supplementation treatment did not influence nematode prevalence, infection status, intensity, or intensity super-shedder status at the population or individual scale. However, food supplementation did significantly reduce the likelihood of individuals becoming abundance super-shedders, a key aspect of parasite infection that has not received previous theoretical attention. Investigating infection heterogeneity, rather than only likelihood of infection and infection intensity, can reveal how anthropogenic changes to food availability can alter infection dynamics. Targeted resource provisioning could help limit transmission potential and mitigate the impact of parasitism without direct medical intervention. We also identified that nematode infection interacts with host age to influence fitness, with infection intensifying age-related trends in survival and reproduction. These findings highlight important patterns in parasite dynamics and host fitness. Food availability may play a role in altering extreme nematode infections such as reducing the likelihood of super-shedding.

Additionally, nematode infections may interact with demographic effects, such as age, in relation to host survival and reproduction. These insights are particularly relevant in systems like bank voles, a key reservoir for Puumala hantavirus, where shifts in parasite transmission and host fitness may have cascading effects on zoonotic disease risk. Therefore, our study carries important management implications for wild populations, suggesting that both nutrition and demographics could be valuable considerations in control strategies.

## Acknowledgements

This work would not have been possible without our collaborators at the Lammi Biological Station and University of Helsinki: John Loehr, Janne Sundell, Esa-Pekka Tuominen, Joni Uusitalo, Tiina Tulonen, Matti Kotakorpi, Riitta Ilola, Jaakko Vainionpää, Tomas Strandin and Tarja Sironen. We thank all students, research station interns, and collaborators who helped collect and process the field data: Alexis Beagle, Anna Bolding, Emilie Bonhomme, Hannah Chan, Chéryline Chanrion, Muriel Chaudhri, Samuel Clague, Juliane Damaschke, Stephanie Du, Lucie Fornilli, Christina Fragel, Mathilde Gaudillère, Shannon Kitchen, Raven Leggett, Teemu Lemola, Eléonore Miston, Nathaniel Mull, Fiorina Muster, Brent Newman, Eunice Oh, Maëlle Pavin De Lafarge, Elizabeth Pellegrini, Austin Rife, Amy Schexnayder, Stephen Shikaze, Aniket Shitole, Anni Simonen, Isabella Stark, Léa Tambareau, and Oliver Xu.

## Author Contributions

**Jasmine S. M. Veitch**: formal analysis, investigation, data curation, writing - original draft, visualization, project administration. **Katherine E. Wearing**: investigation, data curation, writing - review & editing, project administration. **Janine Mistrick**: investigation, data curation, writing - review & editing, project administration. **Meggan E. Craft**: conceptualization, methodology, writing - review & editing, funding acquisition. **Clay E. Cressler**: conceptualization, methodology, writing - review & editing, funding acquisition. **Richard J. Hall**: conceptualization, methodology, writing - review & editing, funding acquisition. **Kristian M. Forbes**: conceptualization, methodology, investigation, writing - review & editing, project administration, funding acquisition. **Sarah A. Budischak**: conceptualization, methodology, investigation, formal analysis, writing - review & editing, supervision, project administration, funding acquisition.

