## Supplemental Information for "Food Supplementation Reduces Nematode Super-Shedding in a Wild Mammal"

### Supplemental Material

Table S1. Generalized linear mixed-effects model for factors associated with population-level nematode prevalence. The parameter 'Occasion' refers to trapping occasion (measure of month of capture) presented as linear term and polynomial term. The parameter 'Year' refers to the year of trapping, comparing the first year of the experiment to the second year (1&2) and the third year (1&3). Bolded terms in table indicate significance (n = 18 months from 14 sites).

| Explanatory Variable | Estimate | SE | P |
|---|---|---|---|
| Food supplementation | -0.076 | 0.045 | 0.091 |
| **Anthelmintic treatment** | **-0.131** | **0.045** | **0.003** |
| **Occasion** | **-1.471** | **0.208** | **<0.001** |
| **Occasion$^2$** | **0.761** | **0.209** | **<0.001** |
| **Year$_{1\&2}$** | **-0.152** | **0.036** | **<0.001** |
| **Year$_{1\&3}$** | **-0.266** | **0.036** | **<0.001** |
| Food supplementation*Anthelmintic treatment | 0.084 | 0.063 | 0.182 |

Table S2. Generalized linear mixed-effects models for factors associated with individual-level nematode infection status and intensity. The parameter 'Pre/post-treatment' distinguishes between an individual's first capture (prior to any anthelmintic treatment effects) and subsequent monthly captures, after the anthelmintic (or sugar water) had the potential to affect nematode infection. The parameter 'Occasion' refers to trapping occasion (measure of month of capture) presented as linear term and polynomial term. The parameter 'Year' refers to the year of trapping, comparing the first year of the experiment to the second year (1&2) and the third year (1&3). Bolded terms in table indicate significance.

| Response Variable | Explanatory Variable | Estimate | SE | P |
|---|---|---|---|---|
| Nematode infection status (n = 2338 captures, 1652 individuals) | Food supplementation$_{Fed}$ | -0.132 | 0.145 | 0.360 |
| | **Anthelmintic treatment$_{Deworm}$** | **-0.365** | **0.175** | **0.037** |
| | Pre/post-treatment | 0.248 | 0.151 | 0.100 |
| | **Occasion** | **-8.933** | **2.984** | **0.003** |
| | **Occasion$^2$** | **11.840** | **2.683** | **<0.001** |
| | **Year$_{1\&2}$** | **-0.934** | **0.118** | **<0.001** |
| | **Year$_{1\&3}$** | **-1.313** | **0.149** | **<0.001** |
| | Sex$_M$ | -0.015 | 0.104 | 0.884 |
| | **Age (head width)** | **0.193** | **0.058** | **<0.001** |
| | **Body condition** | **0.119** | **0.052** | **0.022** |
| | **Population density** | **-0.164** | **0.069** | **0.017** |
| | Food supplementation*Anthelmintic treatment | 0.124 | 0.207 | 0.549 |
| | **Anthelmintic treatment*pre/post-treatment** | **-0.597** | **0.215** | **0.005** |
| Nematode intensity (n = 757 captures, 630 individuals) | Food supplementation$_{Fed}$ | 0.111 | 0.124 | 0.373 |
| | Anthelmintic treatment$_{Deworm}$ | -0.020 | 0.158 | 0.897 |
| | Pre/post-treatment | 0.092 | 0.128 | 0.473 |
| | **Occasion** | **3.902** | **1.720** | **0.023** |
| | **Occasion$^2$** | **3.290** | **1.367** | **0.016** |
| | Year$_{1\&2}$ | 0.060 | 0.104 | 0.564 |
| | Year$_{1\&3}$ | 0.082 | 0.139 | 0.556 |
| | Sex$_M$ | 0.056 | 0.094 | 0.551 |
| | **Age (head width)** | **-0.137** | **0.054** | **0.011** |
| | Body condition | -0.068 | 0.047 | 0.151 |
| | **Population density** | **-0.160** | **0.063** | **0.011** |
| | Food supplementation*Anthelmintic treatment | -0.091 | 0.187 | 0.628 |
| | Anthelmintic treatment*pre/post-treatment | -0.363 | 0.195 | 0.063 |

Table S3. Generalized linear mixed-effects model for effects of experimental treatments and interaction between treatments on likelihood of abundance and intensity super-shedding status, the top 20% of nematode EPG values for all individuals (abundance) or for infected individuals only (intensity). Bolded terms in table indicate significance.

| Response Variable | Explanatory Variable | Estimate | SE | P |
|---|---|---|---|---|
| Abundance super-shedder status (all individuals, n = 2,524 captures, 1,707 individuals) | **Food supplementation$_{Fed}$** | **-0.330** | **0.143** | **0.021** |
| | **Anthelmintic treatment$_{Deworm}$** | **-0.769** | **0.172** | **<0.001** |
| | Food supplementation* Anthelmintic treatment | 0.270 | 0.224 | 0.228 |
| Intensity super-shedder status (infected individuals only, n = 817 captures, 659 individuals) | Food supplementation$_{Fed}$ | 0.394 | 0.259 | 0.128 |
| | **Anthelmintic treatment$_{Deworm}$** | -0.148 | 0.325 | 0.648 |
| | Food supplementation* Anthelmintic treatment | -0.379 | 0.423 | 0.228 |

Table S4. Generalized linear mixed-effects models for factors associated with apparent survival and reproductive status of bank voles. The parameter ‘Pre/post-treatment’ distinguishes between an individual's first capture (prior to any anthelmintic treatment effects) and subsequent monthly captures, after the anthelmintic (or sugar water) had the potential to affect nematode infection. Bolded terms in table indicate significance.

| Response Variable | Explanatory Variable | Estimate | SE | P |
|---|---|---|---|---|
| Apparent survival (n = 1244 captures, 947 individuals) | Anthelmintic treatment$_{Deworm}$ | -0.228 | 0.159 | 0.159 |
| | **Pre/post-treatment** | **-0.635** | **0.162** | **0.010** |
| | Nematode infection status | -0.120 | 0.073 | 0.100 |
| | **Age (head width)** | **0.256** | **0.080** | **0.001** |
| | **Sex$_M$** | **-0.336** | **0.141** | **0.017** |
| | **Reproductive status** | **0.720** | **0.168** | **<0.001** |
| | **Body condition** | **0.217** | **0.076** | **0.004** |
| | Population density | 0.038 | 0.069 | 0.587 |
| | Anthelmintic treatment*pre/post-treatment | 0.035 | 0.296 | 0.905 |
| | **Nematode infection status*age (head width)** | **0.189** | **0.070** | **0.007** |
| Reproductive status (n = 2446 captures, 1676 individuals) | Anthelmintic treatment$_{Deworm}$ | -0.106 | 0.142 | 0.455 |
| | Pre/post-treatment | -0.008 | 0.173 | 0.963 |
| | **Age (head width)** | **1.571** | **0.110** | **<0.001** |
| | Nematode infection status | 0.002 | 0.129 | 0.989 |
| | Anthelmintic treatment*pre/post-treatment | -0.068 | 0.244 | 0.779 |
| | **Nematode infection status*age (head width)** | **0.446** | **0.165** | **0.007** |

Table S5. Generalized linear models for factors associated with individual-level nematode infection status and intensity from the first two months of experiment. The parameter 'Occasion' refers to trapping occasion (measure of month of capture). Bolded terms in table indicate significance.

| Response Variable | Explanatory Variable | Estimate | SE | P |
|---|---|---|---|---|
| Nematode infection status (n = 81 individuals) | Food supplementation | -0.751 | 0.776 | 0.333 |
| | Anthelmintic treatment | -0.388 | 0.832 | 0.641 |
| | Occasion | -0.695 | 0.814 | 0.393 |
| | $Sex_M$ | -0.393 | 0.545 | 0.470 |
| | **Age (head width)** | **0.924** | **0.316** | **0.003** |
| | Body condition | 0.189 | 0.264 | 0.475 |
| | Population density | 0.134 | 0.326 | 0.682 |
| | Food supplementation*Anthelmintic treatment | 0.438 | 1.055 | 0.678 |
| Nematode intensity (n = 58 individuals) | Food supplementation | -0.035 | 0.136 | 0.795 |
| | Anthelmintic treatment | 0.036 | 0.142 | 0.800 |
| | **Occasion** | **0.294** | **0.138** | **0.034** |
| | **$Sex_M$** | **0.259** | **0.109** | **0.017** |
| | Age (head width) | -0.048 | 0.055 | 0.379 |
| | Body condition | -0.033 | 0.056 | 0.557 |
| | Population density | -0.059 | 0.061 | 0.337 |
| | Food supplementation*Anthelmintic treatment | -0.080 | 0.191 | 0.674 |

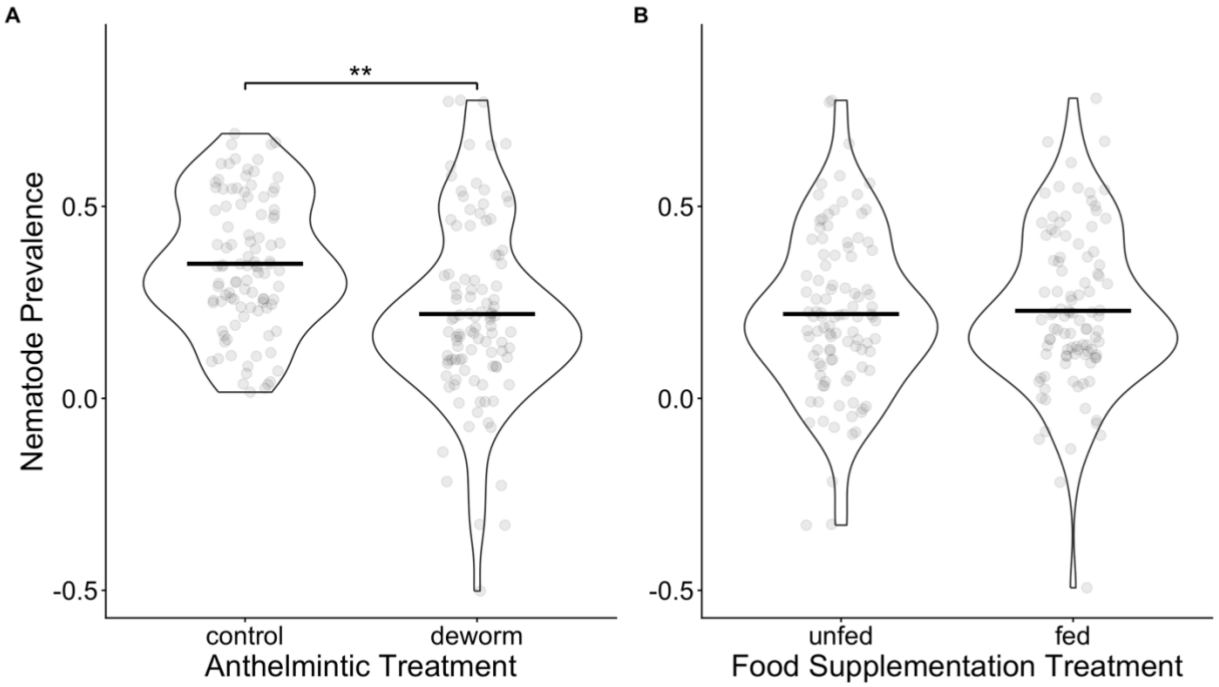


Figure S1. Violin plots displaying the effects of (A) Anthelmintic treatment and (B) food supplementation on nematode prevalence (n = 18 months across 14 sites). Black bars show the group means, and points are jittered for visibility. Points in both panels represent partial residuals. Asterisks indicate significant differences between groups. Anthelmintic treatment, but not food supplementation, reduced nematode prevalence in vole populations.

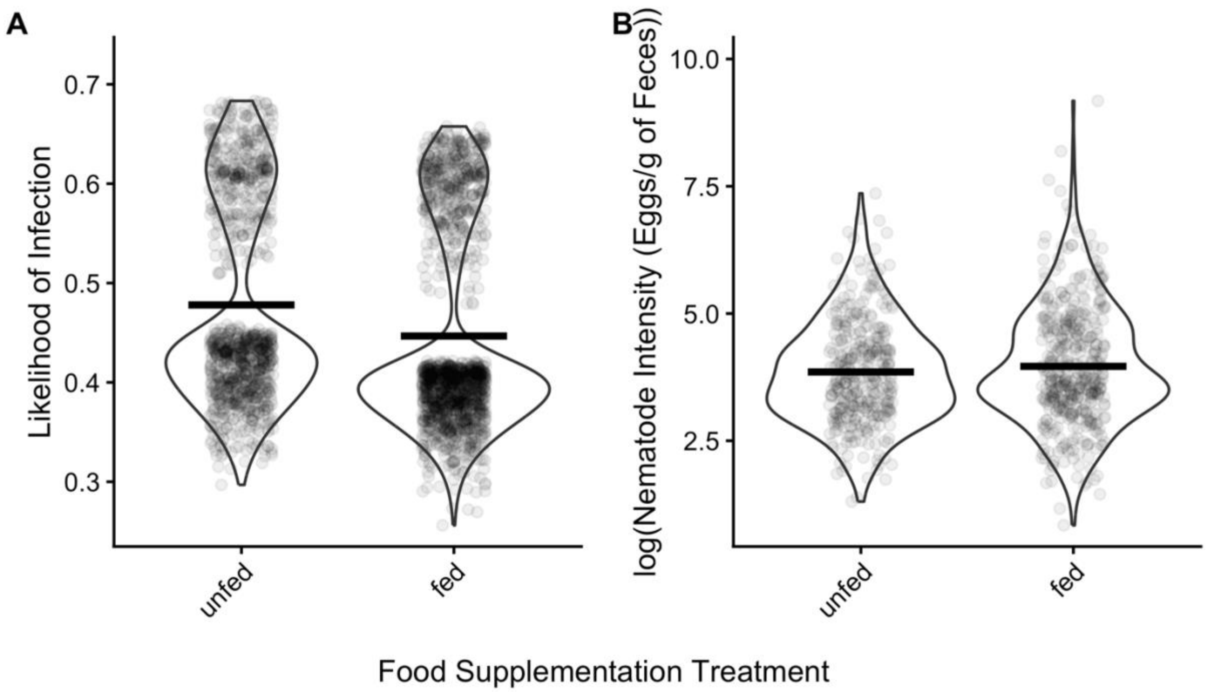


Figure S2. Violin plots displaying the effect of food supplementation treatment on (A) nematode infection status (n = 2338 captures, 1652 individuals) and (B) intensity (n = 757 captures, 630 individuals). Nematode infection status is displayed as the likelihood of infection across food supplementation treatment groups. Nematode intensity includes black bars show the group means and points jittered for visibility. Points in both panels represent partial residuals. Food supplementation had no effect on likelihood of infection or infection intensity at the individual level.

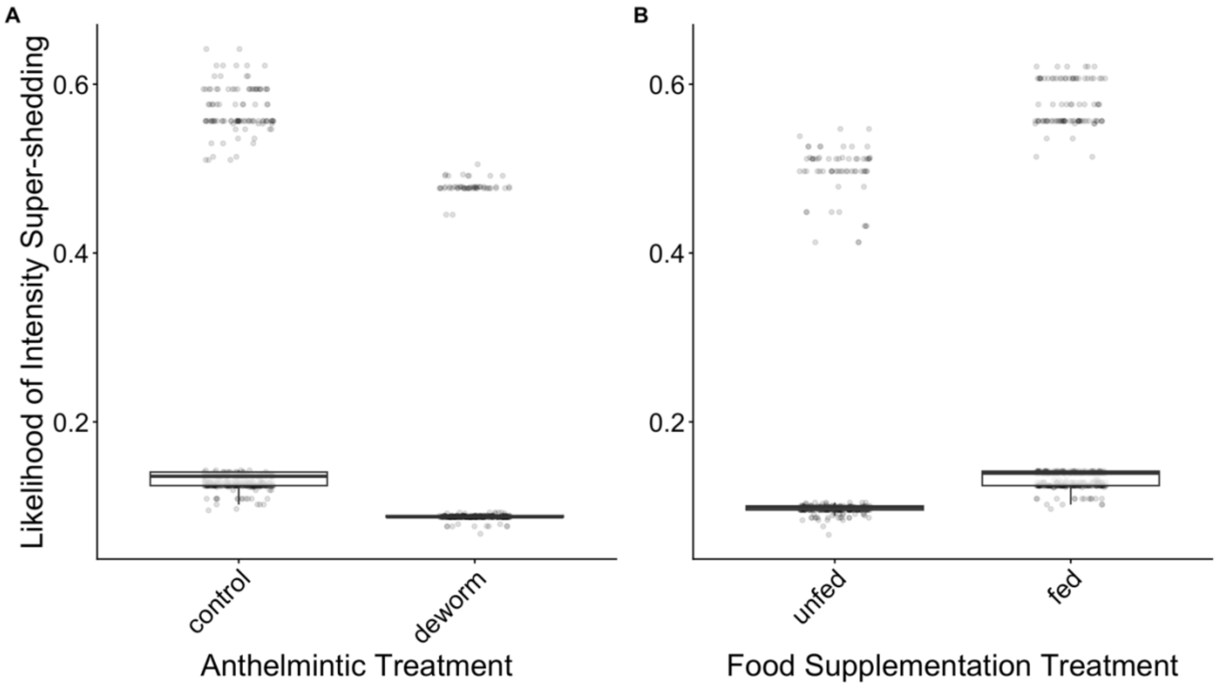


Figure S3. Box plots displaying the effect of (A) Anthelmintic treatment and (B) food supplementation treatment on the likelihood of nematode egg intensity super-shedding (top 20% of nematode EPG values from infected captures; n = 817 captures, 659 individuals). Points in both panels represent partial residuals. Asterisks indicate significant differences between groups. Anthelmintic treatment and food supplementation both reduced the likelihood of an individual becoming a nematode egg super-shedder.

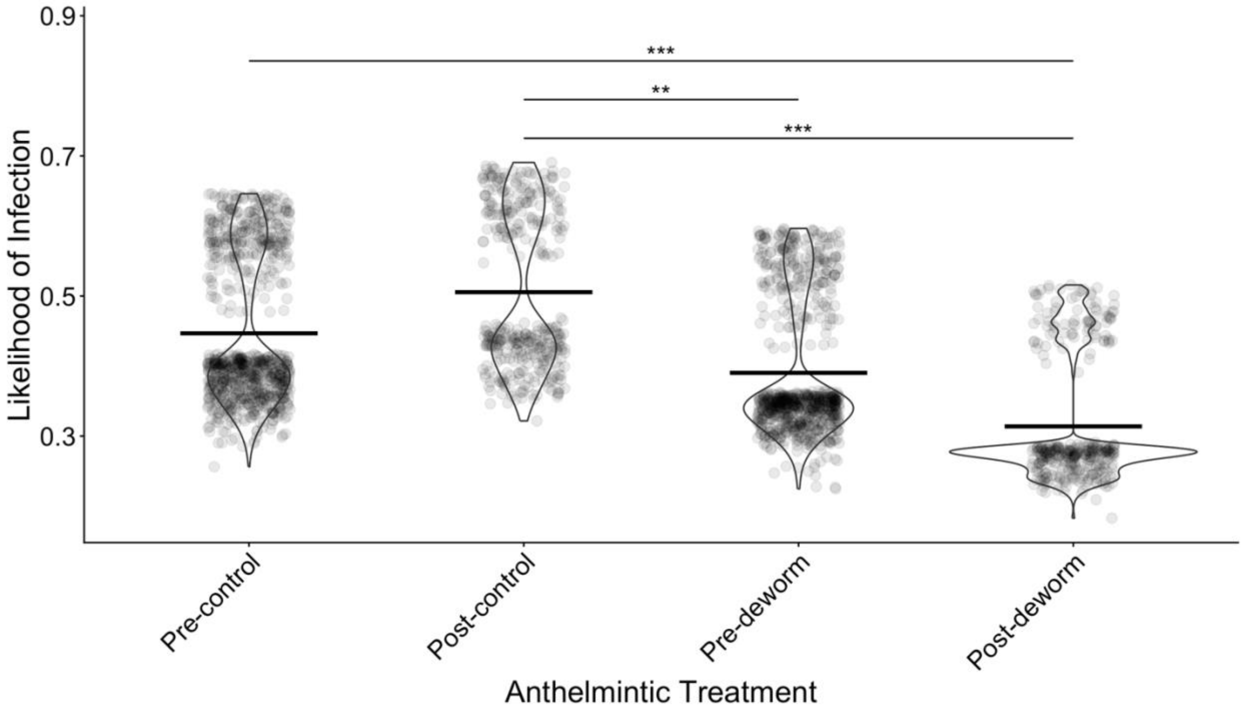


Figure S4. Violin plots displaying the effects of Anthelmintic treatment (A) pre- and (B) post-treatment on nematode infection status (0 = uninfected, 1 = infected; n = 2338 captures, 1652 individuals). Black bars show the group means. Asterisks indicate significant differences between groups. For post-treatment individuals, likelihood of infection was reduced for those in the deworm group compared to those in the control group.

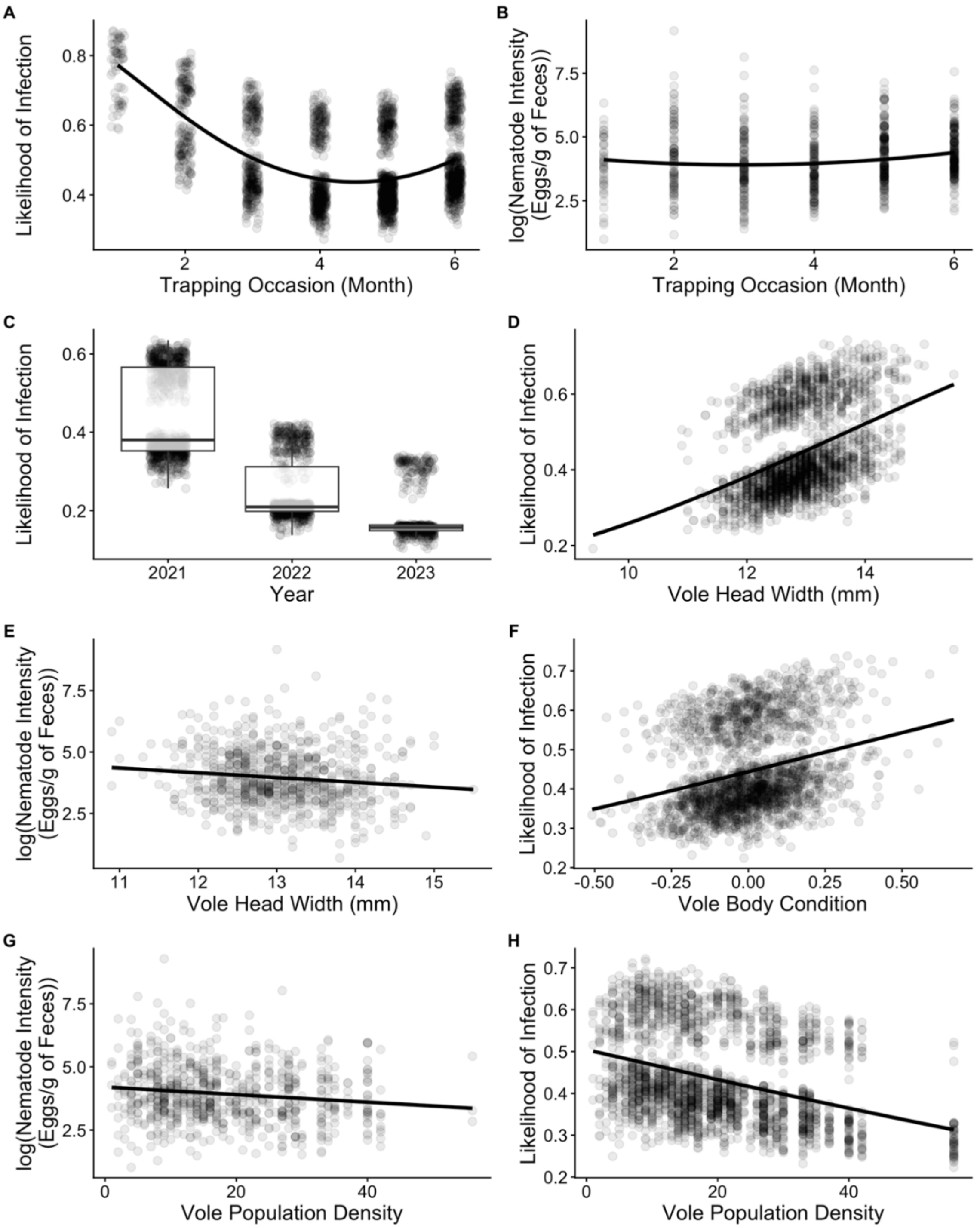


Figure S5. Fitted line and box plots illustrate the relationship between likelihood of nematode infection (n = 2,338 captures, 1,652 individuals) and (A) trapping occasion, (C) year, (D) vole head width (used as a proxy for age), (F) body condition, and (G) population density. The relationship between infection intensity (n = 757 captures, 630 individuals) is shown for (B) trapping occasion, (E) vole head width, and (H) population density. Lines show fitted values from model in Table S2. Points represent partial residuals. For plots displaying likelihood of infection, points are jittered for visibility.
